# Differences in tropical vs. temperate diversity in arthropod predators provide insights into causes of latitudinal gradients of species diversity

**DOI:** 10.1101/283499

**Authors:** Kaïna Privet, Julien Petillon

**Author notes:** Kaïna Privet; Julien Pétillon.

## Abstract

High diversity in tropical compared to temperate regions has long intrigued ecologists. Terrestrial arthropods are among the most speciose orders in tropical rainforests. Previous studies show that arthropod herbivores account for much tropical diversity, yet differences in diversity of arthropod predators between tropical and temperate systems have not been quantified. Here, we present the first standardized tropical-temperate forest comparison of species richness and evenness for understory spiders, a dominant and mega-diverse taxa of generalist predators. Species richness was 13-82 times higher in tropical vs. temperate forests. Evenness was also higher with tropical assemblages having 12-55 times more common and 10-40 times more dominant species. By contrast, proportion of rare species were only up to two times greater than that of temperate measurements. These differences in diversity far surpass previous estimates, and exceed tropical-temperate difference for herbivorous taxa. Thus, the extreme diversity of arthropod predators is associated not only with the higher diversity of prey in tropical vs. temperate ecosystems, but probably also with increased diet breadth of understory spiders in the tropics. This work contradicts the widely accepted hypothesis that tropical diversity is associated with more specialization of predators.

## Introduction

The latitudinal gradient of diversity, that is increase in species richness with decreasing latitude has long been recognized by the scientific community (Pianka 1966). Gradients of diversity in various arthropod taxa from tropical to temperate and even polar ecosystems are well documented through meta-analyses (Willig et al. 2003, but see Hillebrand 2004). Arthropods were particularly studied in tropical rainforests (the species richest terrestrial ecosystem: (Miller et al. 2002) where nearly 1.5 million species are currently described out of an estimated number of 7 million (Hamilton et al. 2010, Stork 2017) or even estimates to 30 million (Erwin 1982). Herbivorous arthropod assemblages have been extensively studied in both tropical and temperate forests with studies of diversity, species richness per plant, host specificity and herbivory pressure. Herbivore arthropod diversity, as well as rate of herbivory, are considered higher in tropical systems compared to temperate counterparts (Lim et al. 2015, Peguero et al. 2017), though evidence of greater host specificity is still controversial (Novotny 2006, Peguero et al. 2017). Such gradients in herbivory diversity can be explained by underlying plant diversity, herbivore diet specialization and plant defense. Less well studied is the possible role of herbivores natural enemies (i.e. predators and parasitoids) on herbivore arthropod diversity (Björkman et al. 2011). Latitudinal gradients in the diversity of omnivore arthropods has also been studied, mainly in ants for which assemblages are clearly species richer in tropical versus temperate systems (Jeanne 1979, Jaffre et al. 2007). For example, canopy assemblages of ants from tropical forests are estimated to be 4 times higher than those from temperate forests (Jaffre et al. 2007). Although ants are considered the main predatory arthropods in tropical rainforests (Floren et al. 2002), they complete a large variety of functional roles (Dejean and Corbara 2003), and their diversity thus does not reflect the diversity of predatory arthropod taxa.

Few studies have examined the latitudinal gradient of predatory arthropod diversity. To date, most of the studies focused on predation pressure, e.g. highlighting that predation pressure increases when latitude decreases (Andrew and Hughes 2005, Novotny 2006, Rodríguez-Castañeda 2013) and sometimes remains constant (Zhang and Adams 2011, Cardoso et al. 2011). Lacking are studies that directly compare the diversity of tropical vs. temperate for predatory arthropods diversity (Schuldt et al. 2013). Basset *et al.* (2012) estimated by indirect comparison that predatory and parasitoid arthropod diversity should be in the same range than herbivore arthropods, with tropical assemblages 2 to 8.4 times more diverse as temperate forest of the same size (Basset et al. 2012). This difference of diversity has been explained by both plant species richness (Basset et al. 2012) and plant phylogenetic diversity (Dinnage et al. 2012), with the suggestion that diversity of predatory arthropods will also mirror plant diversity. But, to our knowledge no study has quantified the ratio of predatory arthropod diversity between tropical and temperate assemblages by direct and standardized comparison (Whitehouse et al. 2009, Schuldt et al. 2013).

Among predatory arthropods, spiders are unique in that they constitute one of the few taxa, if any other, that are exclusively predatory (Birkhofer and Wolters 2012, Pekár and Toft 2015). Only one species of spider is actually known to be exclusively phytophageous (Meehan et al. 2009, Nyffeler et al. 2016) among more than 47,000 species described to date (World Spider Catalog 2018). Spiders are also the seventh most diverse order of animals and among the most abundant predators in terrestrial ecosystems worldwide (Schuldt et al. 2011, Nyffeler and Birkhofer 2017). Spiders thus have an important impact on invertebrate herbivore populations (Schuldt et al. 2011, Nyffeler and Birkhofer 2017) and are suspected to be the main predators of insects (Selden 2016). Moreover, spiders are abundant, have high species richness and occupy a large range of niches in the forest that make them relatively easy to collect in a reasonable amount of time and money (Coddington et al. 1991, Gardner et al. 2008). Due to these characteristics, spiders were proposed as a model group for uncovering large-scale ecological patterns (Cardoso et al. 2011, Birkhofer and Wolters 2012, Malumbres-Olarte et al. 2018). However few studies focused on comparing tropical and temperate spider diversity at large scale (Whitehouse et al. 2009, Schuldt et al. 2013) and none quantify these differences explicitly.

Here, we present the first standardized tropical-temperate point comparison for vegetation-dwelling predatory arthropod (i.e. spider) diversity using the same spatially-replicated sampling protocol. Our objective is to quantify the differences in spider diversity between tropical and temperate forests. We use species richness but also measures of evenness (Shannon and Simpson indices) as variable responses to get a complete view of diversity, including species abundances, evenness and heterogeneity (Chao et al. 2014). First, if spider diversity is associate with herbivorous arthropod diversity, we predict that tropical forests would exhibit 2-8 times higher spider species richness than temperate forests, as estimated by Basset *et al.* (2012) for all herbivore and non-herbivore arthropods. Second, we expect that the evenness of tropical spider assemblages is greater than temperate counterparts due to the commonness of rarity and the more even distribution with low numbers of individuals per species in tropical systems (Nentwig 1993, Chase and Knight 2013). If so, we expect the ratios of common (emphasized by Shannon index) and dominant species (emphasized by Simpson index) would then be in the same range as the species richness ratio. Further, the proportion of rare species (estimated from evenness) should be higher in tropical assemblages (Coddington et al. 2009). To test these predictions, we developed a short and quasi-optimal protocol sampling spiders in tropical and temperate forests. The comparison of the diversity of tropical and temperate assemblage allowed us to highlight the extreme diversity of tropical arthropod predators and to provide new insights into causes of latitudinal gradients of species diversity.

## Material and methods

### Study site

To partially control for potentially confounding effects of microclimate, site fertility, habitat structure, habitat heterogeneity or isolation on species diversity (for a review see Willis & Whittaker 2002), we replicated tropical and temperate sampling in both space (two replicate of tropical and temperate forests) and time.

The tropical sites were two nature reserves in French Guiana (South America) sharing similar climates: La Trinité Reserve (4°35’20’’N; 53°18’1”O) and Nouragues Reserve (4°04’18”N; 52°43’57”O). Both sites are partially and seasonally flooded rainforests and both were sampled during the rainy season, considered as the period of maximum diversity in tropical forest (Gasnier and Höfer 2001, Vedel and Lalagüe 2013). La Trinité reserve protects around 76,900 hectares of forest where the elevation varies between 89 m and 630 m. The Nouragues reserve protects more than 105,000 hectares of isolated tropical rainforest, and elevation varies between 23 m and 465 m. The vegetation of these reserves is typical of the primary lowland rainforest, with few inclusions of palmetto-swamp forests, liana forests and bamboo forests. La Trinité and Les Nouragues were thereafter called tropical forest 1 and tropical forest 2, respectively. Temperate sites were two mixed forests in Britanny (France): the forest of the military camp of Saint-Cyr-Coëtquidan (47°57’50”N; 2°11’30”O) and the state-owned forest of Rennes (48°11’53”N; 1°33’22”O). Both were sampled in summer, the period estimated to have maximal spider diversity (see Hsieh & Linsenmair 2012). The forest of Saint-Cyr-Coëtquidan is around 2,000 hectares of mixed forest included in the 9,000 hectares’ forest massif of Paimpont where the elevation is about 80-90 m. Thanks to the presence of the military camp, this forest has been protected for 150 years. The national forest of Rennes is a forest plots complex covering 3,000 hectares since the eighteen centuries which is recognized as a natural zone of ecological interested (ZNIEFF) and is partly protected by Natura2000 network (1,730 ha). Only irregular forest older than 90 years was sampled in Rennes forest. In both temperate forests, only deciduous (native) tree assemblages were targeted. The vegetation of these forests is typical of the temperate forests with some shrubby species, small trees and climbing plants. The elevation is about 120 m. Saint-Cyr-Coëtquidan and Rennes were thereafter called temperate forest 1 and temperate forest 2, respectively.

### Sampling design and field work

In each forest, the sampling was replicated by using two complementary standardized sampling methods proven to be highly efficient for vegetation-dwelling spiders (Coddington et al. 1991, Cardoso et al. 2008). Two surface-standardized active sampling methods targeting low understory vegetation were selected: beating and sweep netting. These methods were initially used by day and by night but only day data are presented here; Privet *et al.* (submitted) showed no difference of species richness between day and night assemblages at one of the sampling (tropical) sites. Beating vegetation was used to collect spiders living in the shrub, high herb vegetation, bushes, and small trees (hereafter called sampling method 1). A stout stick was used to hit branches or other vegetation to collect falling specimens on a beating tray placed underneath. These samples were conducted in 9 x 9 m quadrats where the vegetation was beat to a height of 2.5 meters. In each forest, 12 quadrats were conducted by four people (two duos) concurrently (six quadrats per duo). Sweep netting (hereafter called sampling method 2) was carried out in the lower herb layer or shrubby vegetation with a sweep net along 20 meters long and one-meter-wide (arm length plus sweep handle) transects. 12 transects were conducted in each forest by the same two persons. This protocol is quasi-optimal (sensu Malumbres-Olarte *et al.* 2017) and was designed for short and intensive survey. Tropical forest 1 was sampled 3-7 December 2010, tropical forest 2, 6-15 December 2013, temperate forest 1, 15-16 June 2015 and temperate forest 2, 22-23 June 2015.

### Sorting and identification

Samples were preserved in 70% ethanol. Individuals were first sorted and identified to family. Temperate adult spiders were sorted and identified to species. Tropical adult spiders were identified to morpho-species based on morphological traits, mainly by observation of genitalia and habitus at 65x magnification. Because only adults are identifiable at species level, juveniles were excluded of the richness measures for all sites, although they represented ∼76% of all the specimens (N=2,846). All specimens are deposited at the University of Rennes 1 (Rennes, France).

## Statistical analysis

We standardized the comparison between the four forests by using a coverage-based approach (Chao and Jost 2012, Chao et al. 2014). The difference between tropical and temperate spider assemblage’s diversity was evaluated by comparing species richness, Shannon and Simpson’s diversity. These comparisons were made by the method of species rarefaction and extrapolation curves based on sample coverage (Gotelli and Colwell 2011, Chao and Jost 2012, Chao et al. 2014). Extrapolations of species richness were realized using the asymptotic Chao1 estimator (Chao 1984). Analyses were completed using the R-based iNEXT package (Chao et al. 2014, Hsieh et al. 2016) with R Software (R Development Core Team 2015) on summed species abundances over the 12 replicates per method per site. iNEXT function was configured at 40 knots and 200 bootstraps replications (method developed by Chao *et al.* 2014). Diversities were compared at equal sample coverage, that yield a less biased measure of assemblage diversity than of equal sample size as assumed by traditional rarefaction methods (Chao and Jost 2012), allowing us to standardize the comparison of spider assemblage diversity between biomes. The common level of sample coverage, named “base coverage”, was determined following Chao *et al.* (2014). Then, comparisons of sample coverage were conducted on rarefied samples: at 38.8% sample coverage for sampling method 1 and at 60% sample coverage for sampling method 2. 95% confidence intervals (CI) were calculated for the three measures of species diversity (species richness, Shannon and Simpson diversity indices) within overlap of CI used to indicate a significant difference at a level of 5% among the expected diversities (Chao et al. 2014). Ratios were calculated as tropical measures compared to temperate ones. For the three diversity measures (e.g., richness, Shannon and Simpson), we selected the lowest and the highest bounds among the measures obtained for the two sampling methods. Proportion of rare species was calculated from the evenness factor (EF) specifying diversity of order 0 (i.e. species richness) and diversity of order 2 (i.e. Simpson) as 1-EF_0,2_ (Jost 2010).

## Results

Based on rarefaction, the sample coverage is nearly two times higher in temperate forests for the two sampling methods and almost any sample size (i.e. number of individuals; Fig.1 A and B). When comparing samples at the same effective sample size for both methods, sample coverage was about 90% in temperate and between 30% and 53% in tropical forests. Thus, even though the same standardized protocol was used in both biomes, temperate samples are 2 to 3 times more complete than tropical ones. Based on the extrapolation for both sampling methods, when the sample size is doubled, the sample coverage increases by 3 to 7% for temperate forests and by 9 to 16% in tropical ones (Fig.1 A and B).

**Figure 1:**
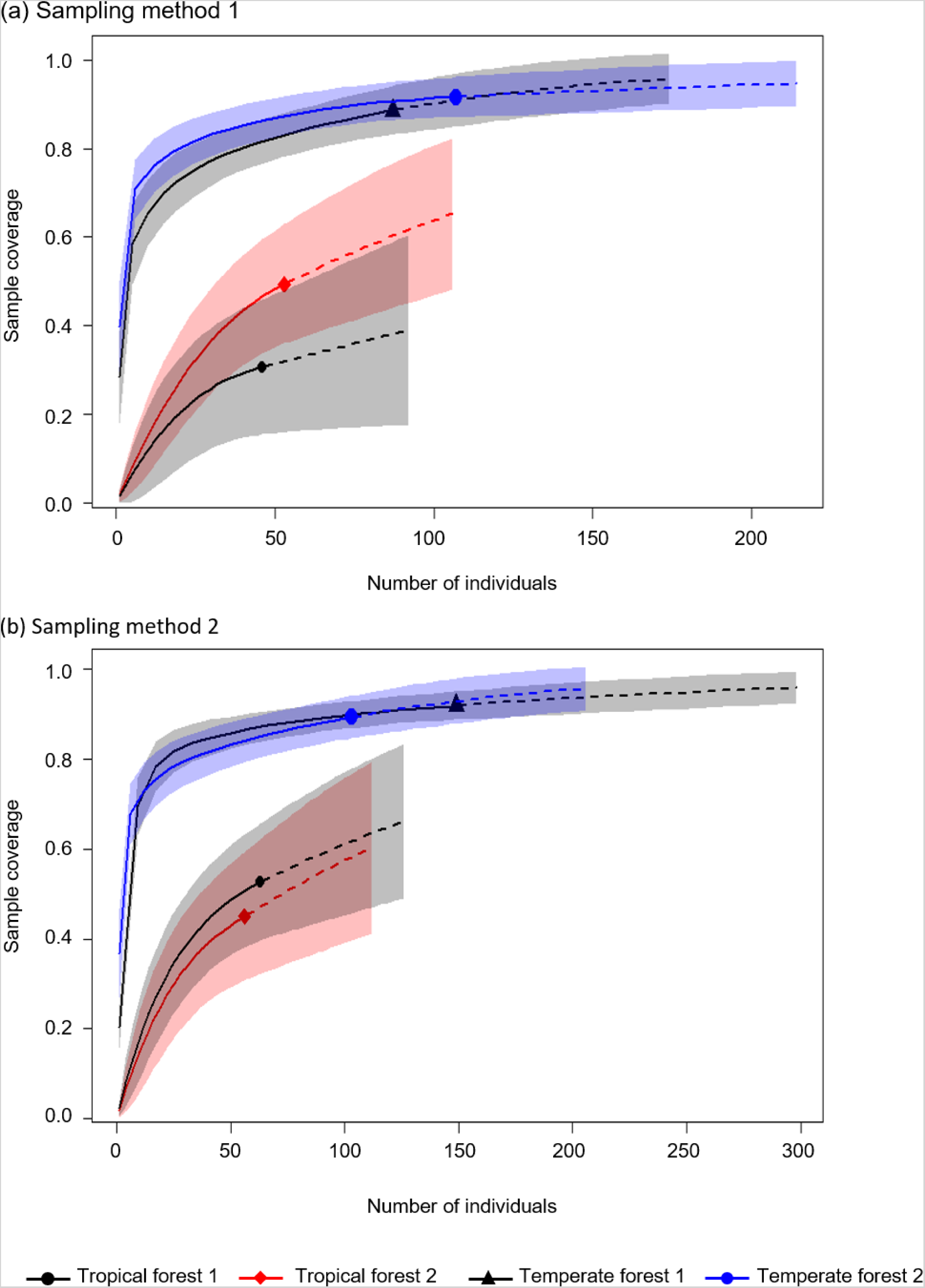
Sample coverage for rarefied samples (solid line) and extrapolated samples (dashed line) as a function of sample size for spider samples collected by (a) sampling method 1 (beating) and (b) sampling method 2 (sweep netting) in in tropical rainforests 1 and 2 (La Trinité and Les Nouragues) and the temperate deciduous forests 1 and 2 (Coëtquidan and Rennes). The 95% confidence intervals are represented in light color and were obtained by a bootstrap method (Chao et al. 2014) based on 200 replications. Reference samples in each forest are denoted by solid markers. For comparison, all curves were extrapolated up to double its reference sample size. The numbers in parentheses are the sample coverage and the number of individuals for reference samples.

When comparing coverage-based diversities of tropical and temperate on reference sample coverages, confidence bands of the replicated sites of tropical and temperate forests do not overlap for either sampling method 1 or for sampling method 2 (Fig. 2 and Fig. 3). Thus, tropical spider assemblages were highly and significantly more diversify than temperate ones for any standardized sample coverage, sampling method and diversity indices used (see detailed results below). Given that sampling methods 1 and 2 consistently showed the same patterns, we will not detail the result per method here.

**Figure 2:**
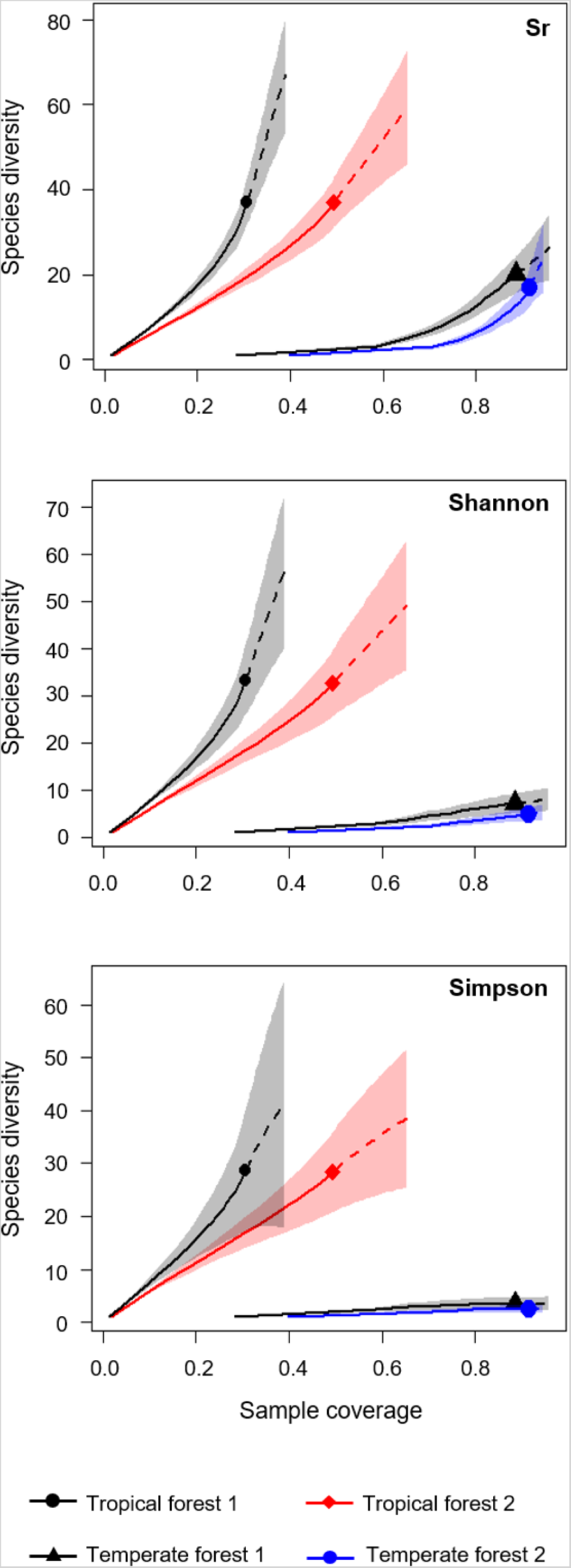
Comparison of the coverage-based rarefaction (solid line) and extrapolation (dashed line) of spider species richness (Sr, left panels), Shannon diversity (middle panels) and Simpson diversity (right panels) collected by sampling method 1 (beating) in tropical rainforests 1 and 2 (La Trinité and Les Nouragues) and the temperate deciduous forests 1 and 2 (Coëtquidan and Rennes). The 95% confidence intervals are represented in light color and were obtained by a bootstrap method (Chao et al. 2014) based on 200 replications. Reference samples in each forest are denoted by solid markers. For comparison, all curves were extrapolated up to double its reference sample size. The numbers in parentheses denote the sample coverage and the observed diversity indices (species richness, Shannon or Simpson) for each reference sample.

**Figure 3:**
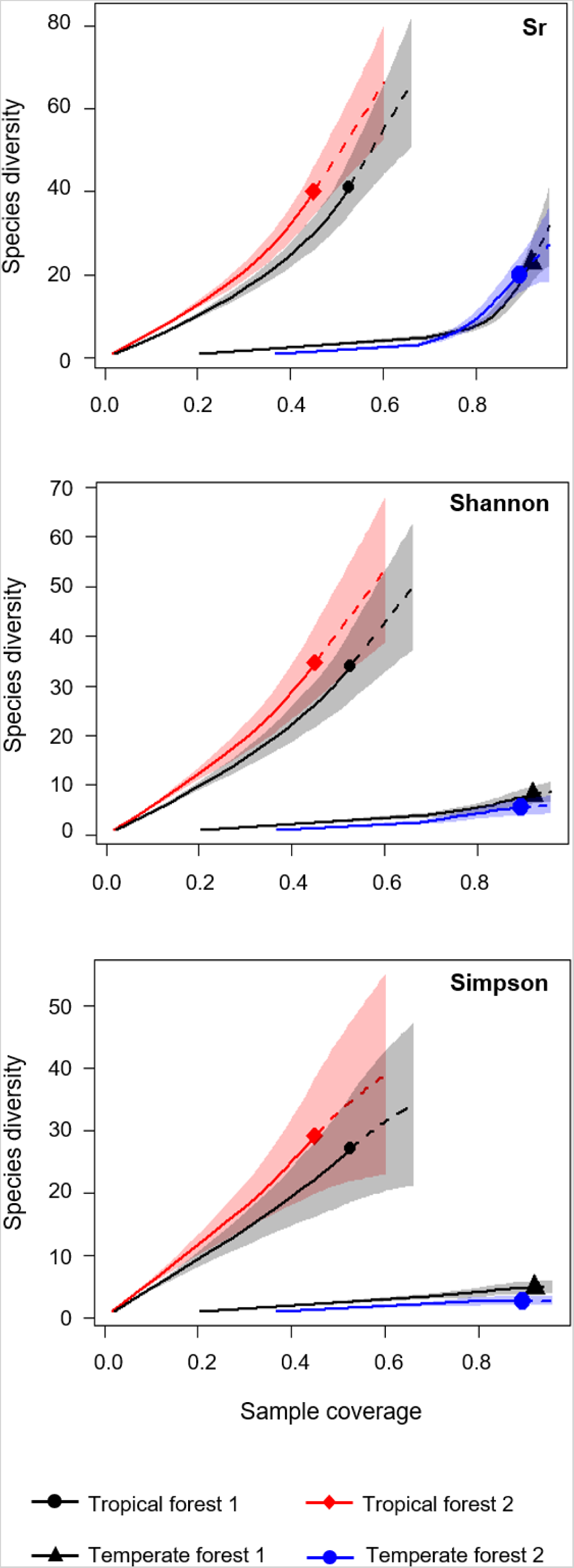
Comparison of the coverage-based rarefaction (solid line) and extrapolation (dashed line) of spider species richness (Sr, left panels), Shannon diversity (middle panels) and Simpson diversity (right panels) collected by sweep netting method in tropical rainforests 1 and 2 (La Trinité and Les Nouragues) and the temperate deciduous forests 1 and 2 (Coëtquidan and Rennes). 95% confidence intervals were obtained by a bootstrap method (Chao et al. 2014) based on 200 replications and are represented in light color. Reference samples in each forest are denoted by solid markers. All curves were extrapolated up to double its reference sample size. The numbers in parentheses denote the sample coverage and the observed diversity indices (species richness, Shannon or Simpson) for each reference sample.

For the base coverage, tropical spider assemblages were 12.9 to 81.6 times richer than temperate ones (Fig. 2 and 3). Difference of diversity between biome is also significant for Shannon diversity and for Simpson diversity. Shannon diversity was 11.6 to 54.6 times higher in tropical assemblages than in temperate counterparts and Simpson diversity was 10.4 to 40.4 times higher in tropical assemblages (Fig. 2 and 3). Finally, proportion of rare species for the base coverage was estimated at 38% and 15% in tropical forests 1 and 2 assemblages sampled by sampling method 1 respectively and 28% and 0% in temperate forests 1 and 2 respectively. When proportion of rare species was estimated from sampling method 2 diversities, there were 42% and 41 % rare species in tropical forest 1 and 2 and 28% and 42% in temperate forests 1 and 2 respectively. Hence, proportion of rare species is not always higher in tropical than temperate but when the case, rare species proportions are up to 2 times more abundant.

## Discussion

Direct and standardized comparison of tropical and temperate predatory arthropod species assemblages’ is a challenging task. Here, we present the first quantification in the difference of diversity between tropical and temperate forest predatory arthropods using understory vegetation spider assemblages, (i) based on a standardized sampling protocol used in among climate and latitudes, (ii) using species richness and diversity indices, and (iii) comparing these diversity parameters by using a standardized statistical rarefaction approach that consider the bias of sample size and species abundance.

Because we used rarefaction to determine richness as well as common and dominant species diversity at a given same level of sample coverage, the resulting measures are not estimates of actual assemblages’ parameters but descriptive measures that allow us to compare tropical and temperate assemblages. Our comparison showed that, with the same level of sample coverage, species richness of tropical spider forest is 13 to 82 times higher that temperate species richness. This magnitude of difference is much greater than expected. Indeed, in previous estimates by Basset *et al.* (2012), tropical arthropod forest diversity is 2 to 8 times higher than in temperate forest of the same size. Several factors can explain the difference in ratios between Basset *et al.* (2012) and this study. First, Basset *et al.* (2012) did not directly compare tropical and temperate diversities, but used data from different studies to determine this ratio and, as acknowledged by the authors, the sampling effort was different in the two biomes. Second, Basset *et al.* (2012) sampled spiders from all strata using 14 different protocols targeting soil, litter, understory, mid-canopy and upper canopy habitats. Thus, the huge difference of diversity we present here between tropical and temperate understory vegetation-dwelling spiders was likely overwhelmed in Basset *et al.* (2012) data by lower diversities from other strata. Third, we used different methods to estimate richness. Basset *et al.* (2012) used six different models to estimate tropical species richness including Chao2 estimator and species accumulation curves based on sampling effort. However, although they used a large range of estimation methods, they did not consider that species richness varies with sample coverage (Chao et al. 2014).

The comparison of evenness showed that tropical species assemblages are up to 55 times greater in tropical than in temperate forests. This is due to a more regular distribution of species illustrated by 12 to 55 times more common species and 10 to 40 times more dominant species. Hence, there are lower numbers of individuals per species in tropical assemblages. The proportion of rare species in tropical spider assemblages was also higher than in temperate assemblages, but only by a factor of two. Thus, the difference of diversity between tropical and temperate spider assemblages is not mainly related to differences in distribution of rare species, but to more even assemblages with lower numbers of individuals per species in tropical assemblages. Weighted measures of diversity (i.e., species evenness and species dominance) are known to provide more comprehensive views of latitudinal gradients (Willig et al. 2003), and actually responded in a way similar to species richness in this study. These results are consistent with our second prediction. Thus, the ratio of spider diversity we measured using a standardized and direct way is up to 30 times what was previously proposed for predators through indirect comparisons.

Our result prompt a reanalysis of the relationship between predatory arthropod diversity, plant species richness and phylogenetic diversity. Basset *et al.* (2012) and Dinnage *et al.* (2012) demonstrated that herbivore and non-herbivore arthropod (including predator) species richness are closely related to plant species richness and phylogenetic diversity by a one-to-one relationship. It was previously estimated that there are 5 to 10 times more plant species per 10,000 km2 and 6 times more tree species per hectare in tropical compared to temperate areas (Barthlott et al. 1996). Hence, the 13-82 times higher species richness of tropical rainforest is vastly higher than what is predicted by plant diversity. Thus, spiders would be 1.2 to 16 times proportionally richer than plants in tropical compared to temperate systems. Although we did not directly measure plant diversity, these results suggest that the relationship between spider and plant diversity in tropical forest is not one-to-one as it was estimated for herbivore and non-herbivore (including spiders) arthropods (Dinnage et al. 2012, Basset et al. 2012).

We suggest that the ratio between plant and spider diversity in tropical forests compared to temperate forests is higher due to difference in diet of spiders in tropical and temperate forests. While most spiders are considered generalist predators (Pekár and Toft 2015), Kozlov *et al.* (2015) highlighted that the latitudinal gradient of diversity is stronger for euryphagous (i.e., non-specialized) spider species than for more specialized ones. Birkhofer & Wolters (2012) also showed that spider diet breadth is higher in tropical environments, and likely associated with net primary production. Together, these observations support the idea that spiders are more diverse in tropical compared to temperate habitats because they feed on more prey species. Thus, the extreme diversity of arthropod predators is associated not only with the higher abundance and diversity of prey in tropical vs. temperate ecosystems, but also increased diet breadth. This suggest the strengths of interactions are greater in tropical regions, not because of increased specialization but because spiders are increasing diet breath (e.g., predation on more prey, but also competition among spiders). Thus, this work contradicts the widely accepted hypothesis that tropical diversity is associated with more specialization (for another example on tropical spiders see Pétillon *et al.* 2018). However, our finding is consistent with Araujo and Costa-Pereira’s study (2013) which focused on the diet breadth of 76 species of vertebrate and invertebrate animal taxa, including 7 predatory arthropods (but not arachnids). They found a higher intraspecific niche variation in tropical regions in connection with the higher diversity of resources, and suggested that the greater within-population niche variation facilitates speciation in tropics. Such a pattern supports the ‘individual variation’ theory suggested as an alternative to ‘niche’ and ‘neutral’ theories of biodiversity. The ‘individual variation’ theory proposed that individual variation allow species to exist to then promote their diversity (Clark et al. 2007, Clark 2010, Violle et al. 2012).

The knowledge brought by our study raises a lot of questions that should be explored in the future. Are top arthropods predators, as spiders, controlling arthropod herbivorous arthropods in the tropics? If so, how can predators be so diverse? Is the diversity of predators increasing much faster than the diversity of prey? How is it that predators can have such remarkable diet breadth, and still coexist (see also Pekár & Toft 2015)? Are predators overlapping in their use of prey more in tropical systems than temperate, or are they somehow specializing in new ways to feed on the same prey?

In conclusion, we highlight a difference of arthropod predator assemblage diversity in tropical vs. temperate forest that far surpass previous estimates, and exceed tropical-temperate difference for herbivorous taxa as well as plant species richness. Thus, the extreme diversity of arthropod predators is associated not only with the higher diversity of prey in tropical vs. temperate ecosystems, but also increased diet breadth which contradict the widely accepted hypothesis that tropical diversity is associated with more specialization but support the individual variation theory.

## Declarations

## Acknowledgements

We are grateful to Frédéric Ysnel who contributed conceiving the project and led the sampling. We thank Coralie Bossu, Alain Canard, Cyril Courtial, Maxime Cobigo, Jennifer Devillechabrolle, Pierre Devogel, El Aziz Djoudi, Audrey Fabarez, Stéphane Icho, Boris Leroy and Vincent Vedel for help during field work, and Marguerite Delaval (Réserves Naturelles de la Trinité and Les Nouragues, French Guiana) for continuous support. We are grateful to George Roderick for the valuable remarks, suggestion regarding this manuscript and English corrections. We thank Rosemary Gillespie for stimulating discussions regarding the results presented here, and Brian Silliman for useful comments.

## Statement of authorship

JP conceived the project, secured the funding and led the sampling. KP sorted, identified spiders and performed statistical analyses. KP wrote the first draft of the manuscript under JP guidance. Both authors discussed the analysis and the results, and contributed to editing the manuscript.

## Data accessibility

Data will be made available on Figshare upon final article acceptance.

